# Nuclear receptor ligand screening in an iPSC-derived *in vitro* blood-brain barrier model identifies new contributors to leptin transport

**DOI:** 10.1101/2022.05.03.490335

**Authors:** Yajuan Shi, Hyosung Kim, Catherine A. Hamann, Elizabeth M. Rhea, Jonathan M. Brunger, Ethan S. Lippmann

## Abstract

**Background:** The peptide hormone leptin exerts its function in the brain to reduce food intake and increase energy expenditure to prevent obesity. However, most obese subjects reflect the resistance to leptin even with elevated serum leptin. Considering that leptin must cross the blood-brain barrier (BBB) in several regions to enter the brain parenchyma, altered leptin transport through the BBB might play an important role in leptin resistance and other biological conditions. Here, we report the use of a human induced pluripotent stem cell (iPSC)-derived BBB model to explore mechanisms that influence leptin transport.

**Methods:** iPSCs were differentiated into brain microvascular endothelial cell (BMEC)-like cells using standard methods. BMEC-like cells were cultured in Transwell filters, treated with ligands from a nuclear receptor agonist library, and assayed for leptin transport using an enzyme-linked immune sorbent assay. RNA sequencing was further used to identify differentially regulated genes and pathways. The role of a select hit in leptin transport was tested with the competitive substrate assay and after gene knockdown using CRISPR techniques.

**Results:** Following a screen of 73 compounds, 17β-estradiol was identified as a compound that could significantly increase leptin transport. RNA sequencing revealed many differentially expressed transmembrane transporters after 17β-estradiol treatment. Of these, cationic amino acid transporter-1 (CAT-1, encoded by SLC7A1) was selected for follow-up analyses due to its high and selective expression in BMECs *in vivo*. Treatment of BMEC-like cells with CAT-1 substrates, as well as knockdown of CAT-1 expression via CRISPR-mediated epigenome editing, yielded significant increases in leptin transport.

**Conclusions:** A major female sex hormone, as well as an amino acid transporter, were revealed as regulators of leptin BBB transport in the iPSC-derived BBB model. Outcomes from this work provide insights into regulation of peptide hormone transport across the BBB.

## Introduction

The brain harmonizes energy homeostasis by interacting with peripheral tissues. Leptin, a 16 kDa peptide hormone secreted by adipocytes, is released into the blood circulation and crosses the blood-brain barrier (BBB) in several regions of the brain to reduce food intake and increase energy expenditure [1]. However, despite serum leptin levels being elevated in direct proportion to the amount of body fat, the efficacy of its anorexic effect is decreased in obesity [2]. Most obese subjects exhibit resistance to leptin, and this resistance impedes sustained weight loss [3]. Intriguingly, mouse models of obesity are still responsive to intracerebroventricularly delivered leptin with high sensitivity compared to peripheral infusion, indicating that impaired brain access via diminished BBB transport may contribute to leptin resistance and other biological conditions [4, 5].

Unlike the leaky endothelium of other vascular beds in the body, the BBB is a highly selective interface. The BBB, whose functional properties are comprised by the brain microvascular endothelial cells (BMECs) that form the inner layer of the capillary vessel wall, strictly regulates bidirectional molecular exchange between the bloodstream and the brain [6]. Except for the free diffusion of gases and small lipophilic drugs, most molecules are transported to and from the brain through a cohort of specific receptors, transporters, and channels [7]. Similar to other peptide hormones, leptin influx is enabled through saturable receptor-mediated transcytosis at the BBB [8]. Although leptin brain uptake has been noted in regions not shielded by the BBB [8–10], the necessity of leptin BBB penetration has been confirmed in preclinical obesity models [11]. At the BBB, the leptin receptor (ObR) has been identified as a transporter of leptin [12]. However, inhibition [13] or deletion [14] of ObR at the BBB in cellular and *in vivo* models does not fully diminish leptin transport, suggesting that this process may not be ObR-dependent and that other mechanisms exist to regulate leptin uptake in the brain.

*In vivo* studies linking BBB transport to specific genes are challenging to conduct due to low throughput. In contrast, *in vitro* BBB models have higher throughput and are more amenable to screening approaches. However, most *in vitro* models are not able to fully recapitulate the BBB properties. In particular, to study molecular transport, models must possess high trans-endothelial electrical resistance (TEER) that limits passive diffusion. The advent of human induced pluripotent stem cell (iPSC) technology has provided BMEC-like cells that exhibit robust BBB properties [15–20], including a cohort of essential transporters and restrictive paracellular permeability to facilitate central nervous system (CNS) drug-permeability studies [21, 22], investigate disease biology [23–26], and study mechanisms of molecular transport [27–29]. Outcomes from these studies can provide critical insights into BBB biology, particularly when validation work is performed in physiologically relevant *ex vivo* and *in vivo* systems [30].

Here, we aimed to use iPSC-derived BMEC-like cells to better understand transcytosis of leptin across the BBB. Our strategy was to screen a nuclear receptor agonist library and assessed changes to leptin transport. We chose this strategy because nuclear receptors control a broad range of biological processes through activation by small lipophilic ligands [31]. Nuclear receptor signaling pathways have been implicated in nutrient and efflux transporter function in solid organs and tissue barriers such as the gut, peripheral endothelium, and BBB [32–34]. These pathways also play important roles in energy metabolism, lipid dysfunction, and obesity-related diseases, including manipulating the process of lipogenesis and gluconeogenesis, preventing the onset of obesity, and reversing the induced obesity in obese mouse models [35–37]. Moreover, nuclear receptor signaling influences the development, maintenance, and function of the BBB [16,38,39]. From this screen, we determined that treatment of BMEC-like cells with 17β-estradiol yielded a significant increase in leptin transport. RNA sequencing of 17β-estradiol-treated BMEC-like cells revealed a set of differentially regulated genes, and the gene ontology analysis on molecular functions emphasized signaling pathways involving transmembrane transporter activity. For validation, we manipulated one particular hit—cationic amino acid transporter 1 (CAT-1)—through functional and genetic assays and demonstrated changes to leptin transport. Overall, our work provides new insight into peptide hormone transport across the BBB. We expect this study will provide a framework for screening and assessing how specific peptide hormones and nutrients are transported across the BBB.

## Methods

### iPSC Maintenance and Differentiation

CC3 iPSCs were maintained and differentiated at 37°C in a humidified CO_2_ incubator following our previously published methods [18]. In brief, CC3 iPSCs were maintained in E8 medium on 6-well tissue culture plates (Fisher Scientific #07-20083) coated with growth-factor reduced Matrigel (Corning #354230), and cells were passaged every 3-4 days once reaching 70-80% confluency using pre-warmed Versene (Fisher Scientific #15-040-066).

To differentiate iPSCs to BMEC-like cells, CC3 iPSCs were washed once with DPBS (Fisher Scientific #14-190-144), dissociated with Accutase for 3-4 minutes (Fisher Scientific #A1110501), collected via centrifugation, and resuspended into E8 medium supplemented with 10 µM Y27632 (Tocris #1254). Cells were seeded on 6-well tissue culture plates coated with Matrigel at the density of 150,000 cells/well. Twenty-four hours after seeding (noted as Day 0 in Figure 1A), the medium was switched to E6 medium and then changed every 24 hours for 4 days. On Day 4, cells were changed to EC medium consisting of human endothelial serum-free media (hESFM; Thermo Fisher Scientific #11111044), 0.25X B27 supplement (Fisher Scientific #17-504-044), 20 ng/mL basic fibroblast growth factor (bFGF; PeproTech #100-18b), and 10 µM all-trans retinoic acid (RA; Sigma-Aldrich #R2625); no media changes were performed for 48 hours. On Day 5, 24-well Transwell filter inserts (Fisher Scientific #07-200-154) were coated with 55 µL extracellular matrix (ECM) containing 400 μg/mL collagen IV (Sigma-Aldrich #C5533) and 100 μg/mL fibronectin (Sigma-Aldrich #F1141) in UltraPure DNase/RNase-free distilled water (Thermo Fisher Scientific #10977015). On Day 6 (noted as Subculture Day -1 in Figure 1A), differentiated cells were washed with DPBS, dissociated with Accutase for 20-30 minutes, collected via centrifugation, and resuspended in EC medium at the ratio of 1 well to 600 µL EC medium. 100 µL of cell suspension were plated on pre-coated Transwell inserts while 600 µL of EC medium were added to the basolateral chamber. On Day 7, referred to as Day 0 of subculture, the medium was changed to EC medium without bFGF and RA on both the apical and basolateral sides. Select experiments were conducted on Subculture Day 1 (Figure 1A) and respective medium modifications are stated below.

**Figure 1.**
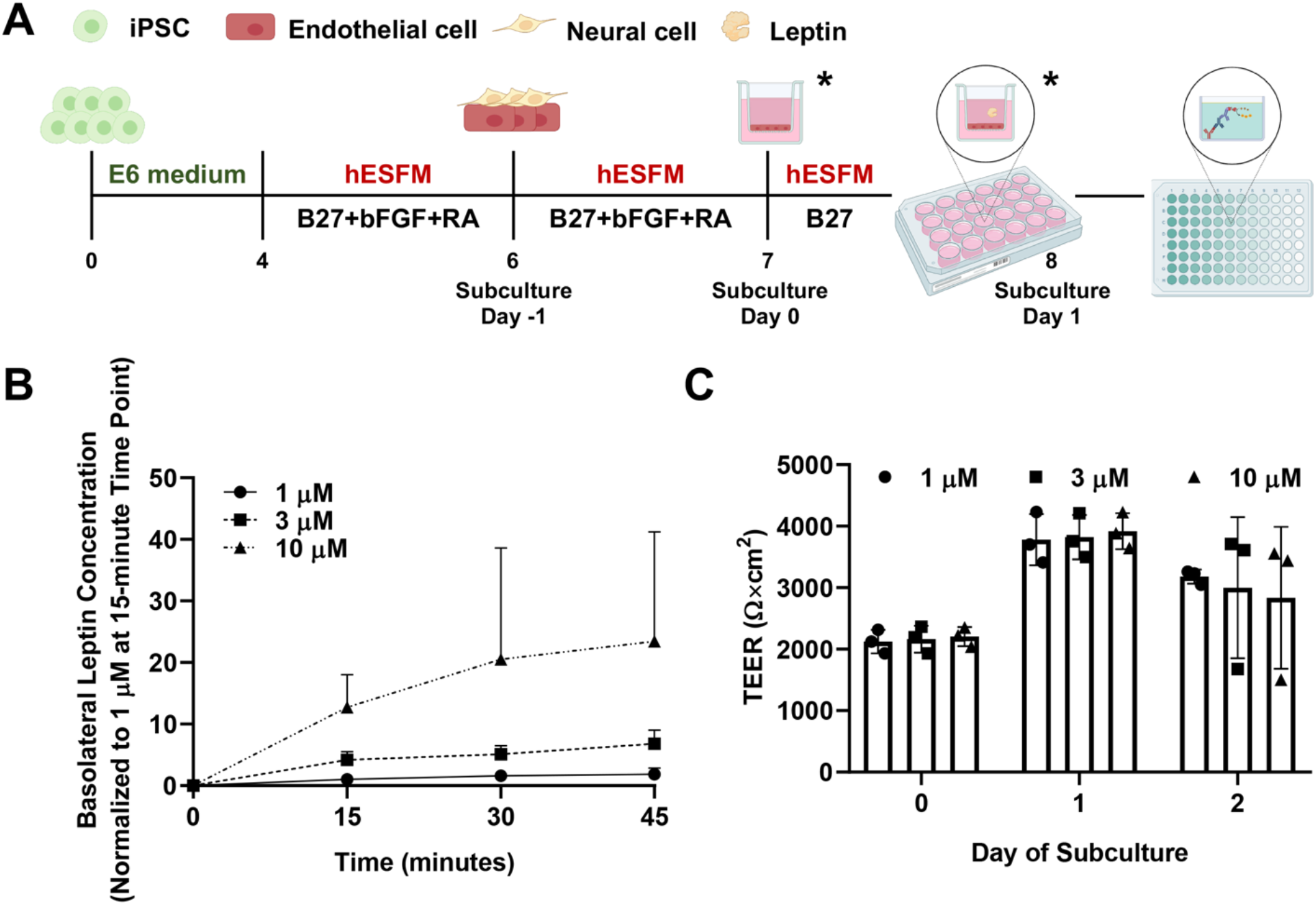
Establishment of leptin transport assay. (A) Experimental timeline for iPSC differentiation and initiation of assays. Leptin transport was always assessed on Day 1 of subculture. Modifications were made on Day 0 or/and 1 of subculture to reflect different experimental setups (time points indicated by asterisks). (B) Temporal leptin transport through BMEC-like cells as a function of leptin concentration in the apical chamber. All samples were measured with ELISA, converted to concentration using a standard curve, and normalized to the 1 µM condition at the 15-minute time point. Data are presented as mean ± standard deviation from n=3 biological replicates. (C) BMEC-like cells demonstrate the high-fidelity passive barrier function before and after the leptin transport assay. TEERs are represented as mean ± standard deviation in a single biological replicate (n=3 measurements from each filter). Trends were confirmed in two additional biological replicates.

### TEER Measurements

After plating cells on Transwell inserts, TEER was measured every 24 hours on Day 0, Day 1, and/or Day 2 of subculture using the EVOM2 voltohmmeter (World Precision Instruments) with STX2 chopstick electrodes (World Precision Instruments). TEER was measured in an empty Transwell containing no cells as the baseline control. Three measurements were taken from one filter at each time point. TEER values were corrected by subtracting the empty filter reading and multiplying by the surface area of the filter (0.33 cm^2^).

### Initial Assessment of Leptin Transport

Recombinant human leptin (R&D Systems #398-LP) was reconstituted at 1 mg/mL in sterile 20 mM Tris-HCl (Fisher Scientific #MT46031CM) at pH of 8.0. On Day 1 of subculture (Figure 1A), TEER was measured to ensure barrier induction in the BMEC-like cells. Leptin was added to hESFM medium with B27 at the concentration of 1 µM, 3 µM, and 10 µM. At the start of this assay, the medium on the apical chamber of the 24-well Transwell insert was aspirated and then replaced with 100 µL of leptin working solution. 100 µL of medium was then collected from each basolateral chamber. At the same time, 100 µL of fresh hESFM medium with B27 were added back to the basolateral chamber to maintain a consistent medium volume. Basolateral medium collection was repeated every 15 minutes for 45-60 minutes total as needed. Each plate was returned to the incubator after medium extraction and replacement. Collected samples were stored at −80°C, and leptin concentration was measured through ELISA as described below.

### Nuclear Receptor Ligand Screening and Validation

To identify possible ligands involved in leptin transport regulation, the SCREEN-WELL Nuclear Receptor ligand library (Enzo Life Sciences #BML-2802-0100), consisting of 73 compounds dissolved in Dimethyl sulfoxide (DMSO), was used in primary ligand screening. As previously described, BMEC-like cells differentiated from iPSCs were purified onto ECM-coated 24-well Transwell filters in EC medium. TEER was measured approximately every 24 hours after subculture to confirm passive barrier function. On Day 0 of subculture, compounds were added at the concentration of 10 µM into EC medium lacking bFGF and RA. Equivalent volume of DMSO (Sigma-Aldrich #D8418) was used as a negative control. Medium on both sides of Transwell inserts were replaced with medium supplemented with respective select compound or DMSO. 24 hours after the medium change, BMEC-like cells received a medium change only in the apical compartment, respectively. This medium was composed of EC medium lacking bFGF and RA, 10 µM of select compound or DMSO, and 3 µM of leptin. After a 30-minute incubation at 37°C, 100 µL of the medium was extracted from the basolateral compartment in each well and transferred into individual microfuge tubes. Collected medium was stored at −80°C and measured using ELISA as described below. During the primary screening, each condition was assayed across duplicate filters. After analyzing data from the primary screen, 7 select ligands and a DMSO control were re-examined in one 24-well plate with triplicate Transwell filters. The leptin transport assay was conducted as described above. This process was repeated across 3 biological replicates to enable statistical assessments and evaluate reproducibility.

### ELISA Assay

Leptin concentration within all collected samples was determined using the Human Leptin Standard ABTS ELISA Development Kit (PeproTech #900-K90) and ABTS ELISA Buffer Kit (PeproTech #900-K00) according to the manufacturer’s suggestions with subtle adjustments.

Prior to the assay, solutions were diluted and adjusted to pH 7.2 using the Orion Star A221 benchtop pH meter (Thermo Fisher Scientific #STARA2110). On the first day, 100 µL of 1 µg/mL capture antibody in 1X PBS was added into each well of one ELISA microplate (Corning #3590). The plate was sealed with polyester adhesive sheet and incubated overnight at room temperature. On the second day, the plate was washed four times using 300 µL of 1X wash buffer and blotted on a paper towel to remove residual buffer after the last wash. 300 µL of block buffer was then added into each well and incubated for 2 hours at room temperature. During this time, collected samples were diluted into 1X Diluent buffer using the optimized dilution factor (1 µM leptin, 1:10; 3 µM leptin, 1:25; 10 µM leptin, 1:60). Human leptin standard was diluted into 1X Diluent at the concentration of 4000 pg/mL, 3000 pg/mL, 2000 pg/mL, 1000 pg/mL, 500 pg/mL, 250 pg/mL, 125 pg/mL, and 0 pg/mL to generate a standard curve. After blocking, plate was washed with wash buffer and filled with 100 µL of standard or diluted sample in each well in duplicates or triplicates as needed. The plate was not disturbed until sample incubation was finished after 2.5 hours at room temperature. Following sample incubation, the plate was washed again. Then, 100 µL of 0.25 ug/mL detection antibody in 1X Diluent was added into each well of the plate to incubate for 2 hours at room temperature. After incubation with the detection antibody, the plate was washed again and replaced with 100 µL of Avidin-HRP conjugate in 1X Diluent at the ratio of 1:2000. The plate was then incubated with Avidin-HRP conjugate for 30 minutes at room temperature and washed again. Following washing, 100 µL of ABTS was added to each well and incubated at room temperature for color development. The ELISA plate was monitored on a BioTek Synergy H1 multi-mode microplate reader at 405 nm with wavelength correction set at 650 nm for approximately 60 minutes at room temperature. A standard curve was then created in GraphPad Prism using known leptin concentrations and used to convert all experimental measurements to concentration.

### RNA Sequencing and Pathway Analysis

Bulk RNA-sequencing was applied to profile transcriptomic changes under 17β-estradiol treatment. BMEC-like cells were purified onto 6-well plates, and 24 hours after subculture, 10 µM of 17β-estradiol or DMSO was prepared in EC medium lacking bFGF and RA and added to the cells. After 24 hours, BMEC-like cells were washed once with DPBS, lifted using a cell lifter (Fisher Scientific #08-100-240), and collected via centrifugation. Resultant cell pellets were lysed in TRIzol reagent (Thermo Fisher Scientific #15596026) at room temperature for 10 minutes and stored at −80°C. Three biological replicates were collected per condition. Following the manufacturer’s guideline, RNA from each sample was purified using a Direct-zol RNA Miniprep kit (Zymo Research #R2051) simultaneously. Purified RNA samples were submitted to the Vanderbilt Technologies for Advanced Genomics core and sequenced on an Illumina NovaSeq6000. Raw sequencing reads were trimmed and aligned to human genome (GRCh38) with HISAT2 2.2.0 [40], and genomic features were obtained using the featureCounts function [41]. Differential gene expression and meta read counts were analyzed with edgeR [42]. Gene ontology enrichments analysis was performed on genes with p<0.05 and fold change greater than 1.5 using WEB-based Gene SeT AnaLysis Toolkit (WebGestalt) [43].

### Amino Acid Treatments

On Day 1 of subculture, the medium in the apical chamber was replaced with 100 µL of EC medium lacking bFGF and RA and containing 3 µM of leptin and 3 mM of amino acid substrate. After 30 minutes, 100 µL of medium was collected from the basolateral chamber and stored at −80°C. ELISA was then used to quantify leptin concentration as previously described. In terms of amino acids, L-Cystine (Sigma-Aldrich #C8755) was selected as the substrate of cystine/glutamate transporter (encoded by *SLC7A11*), L-Serine (Sigma-Aldrich #S4311) was selected as the substrate of neutral amino acid transporter A (encoded by *SLC1A4*) and neutral amino acid transporter B(0) (encoded by *SLC1A5*), and glycine (Sigma-Aldrich #G8790) was selected as the substrate of glycine transporter 1 (encoded by *SLC6A9*). CAT-1 (encoded by *SLC7A1*) and CAT-3 (encoded by *SLC7A3*) had 4 substrates including L-Lysine hydrochloride (Fisher Scientific #BP386), L-Histidine (Sigma-Aldrich #H6034), L-Ornithine monohydrochloride (Sigma-Aldrich #O6503), and L-Arginine (Acros Organics #104991000).

### CRISPR Plasmid Design and Construction

To produce a Sleeping Beauty (SB) transposon vector for inducible expression of the CRISPRoff-expression cassette, the SB transposon from Addgene plasmid 60495 (gifted from Eric Kowarz [44]) was modified to replace the constitutive RPBSA promoter-driven expression of rtTA-advanced with EF-1α-driven expression of TetOn-3G. This transposon vector then served as the backbone for cloning the CRISPRoff-v2.1 protein, which consists of a fusion protein of the catalytic domains from DNMT3A and DNMT3L to dCas9, blue fluorescent protein (BFP), and KRAB, downstream of the tetracycline response element (TRE) promoter. The TetOn-3G vector was digested with SfiI, and a PCR amplicon containing the CRISPRoff-v2.1 protein from Addgene plasmid 167981 (gifted from Luke Gilbert [45]) was inserted into the backbone downstream of the TRE using NEBuilder HiFi DNA Assembly Master Mix (NEB #E2621) following the manufacturer’s recommendations. The final transposon contains the constitutive TetOn-3G with TRE-driven CRISPRoff.

A second SB transposon vector containing the U6-sgRNA expression cassette was produced by cloning two PCR fragments into the backbone of Addgene plasmid 60495. One insert consisted of the U6-sgRNA cassette from Addgene 71236 (gifted from Charles Gersbach [46]). This cassette contains two BbsI cleavage sites that enable cloning of target-specific spacer sequences between the U6 promoter and the gRNA scaffold. The second insert included the murine PGK promoter upstream of the hygromycin B phosphotransferase transgene. The SB transposon vector backbone, including the SB inverted terminal repeats, was PCR amplified, and the three amplicons were assembled using the NEBuilder HiFi DNA Assembly Master Mix following manufacturer’s recommendations. Three single gRNAs (sgRNAs) targeting *SLC7A1* and three non-targeting control sgRNAs were selected from a published CRISPR library [47]. Sequences are available in Table S1.

These transposon vectors were propagated in DH5α competent cells and co-transfected along with a plasmid encoding the hyperactive enzyme SB 100x Transposase (Addgene 34879, gifted from Zsuzsanna Izsvak [48]) as described below.

### iPSC Transfection, Sorting, and Differentiation

CC3 iPSCs were maintained in mTeSR Plus medium (StemCell Technologies #100-0276) before transfection. To generate a stable cell line with inducible expression of CRISPRoff, the TransIT-LT1 reagent (Mirus Bio #MIR2304) was used to deliver the SB 100x transposase and transposon vectors. First, 250 μL of mTeSR Plus medium containing 1 μL of 10 mM Y27632 was added into a well of a Matrigel-coated 12-well plate. Then, a mixture of plasmids encoding SB transposase (1 µg) and CRISPRoff (1 µg) was prepared in 200 µL of Opti-MEM I reduced serum medium (Thermo Fisher Scientific #319850-62) supplemented with 6 µL of TransIT-LT1. After a 15-minute incubation at room temperature, the transfection mix was added into the well of a 12-well plate. CC3 iPSCs were rinsed once with DPBS, dissociated with Accutase for 3-4 minutes, collected via centrifugation, and resuspended into mTeSR Plus medium at the density of 1.5×10^6^ cells/mL. 500 µL of this cell suspension was added into the well containing the transfection mix. 24 hours after transfection, the transfection medium was exchanged for fresh mTeSR Plus medium.

These same iPSCs were transfected a second time using the methods described above. This second time, a mix of plasmids encoding SB transposase (1 µg) and sgRNAs (1 µg total of 3 pooled gRNA vectors) were used. 24 hours after transfection, transfected cells were changed to mTeSR Plus medium supplemented with 115 µg/mL of Hygromycin B (Sigma-Aldrich #H3274). 2 days later, cells were changed to mTeSR Plus medium supplemented with 115 µg/mL of Hygromycin B and 2 µg/mL of doxycycline hyclate (Sigma Aldrich #D9891). The following day, cells were sorted with a BD FACSAria lll based on BFP expression and plated onto Matrigel-coated plates in mTeSR Plus medium supplemented with 10 µM Y27632. After 3 days, sorted iPSCs were changed to E8 medium supplemented with 2 µg/mL of doxycycline and differentiated into BMEC-like cells and utilized for leptin transport experiments as described above.

### Western Blot Analysis

iPSCs were harvested 9 days after the initiation of doxycycline treatment. Cells were lysed in RIPA buffer (Sigma Aldrich #R0278) supplemented with 1% v/v of protease inhibitor cocktail (Sigma-Aldrich #P8340) and 1% v/v of phosphatase inhibitor cocktail 3 (Sigma-Aldrich #P0044). After 30-minute lysis on ice, samples were centrifugated at 12,000xg for 15 minutes at 4°C and the supernatants were collected. Protein concentrations were measured with a Pierce BCA protein assay kit (Thermo Fisher Scientific #23225). 25 µg of total protein was separated on 4-20% Criterion TGX precast midi protein gels (Bio-Rad #5671094) and transferred onto nitrocellulose membranes (Thermo Fisher Scientific #IB23001) using an iBlot 2 gel transfer device (Thermo Fisher Scientific #iB21001). Membranes were blocked with Odyssey blocking buffer (Li-Cor #92750000) at room temperature for 30 minutes, incubated with Odyssey blocking buffer containing primary antibody and 0.05% Tween 20 (Sigma-Aldrich #P9416) at 4°C overnight, and incubated with 1X Tris Buffered Saline (TBS, Corning #46-012-CM) containing respective secondary antibody and 0.05% Tween 20 at room temperature for 2 hours. Membranes were imaged with Odyssey XF (Li-Cor), and band intensity was determined using Image Studio software.

### qPCR Analysis

Total RNA was extracted on Day 1 of subculture and purified using a Direct-zol RNA Miniprep kit as described above. cDNA was generated using a High-Capacity cDNA Reverse Transcription kit (Applied Biosystems #4368814). qPCR was performed in a 20 µL of reaction mix containing 9 µL of cDNA template, 10 µL of TaqMan Universal PCR Master Mix (Applied Biosystems #4304437), and 1 µL of TaqMan probe (Thermo Fisher Scientific, *SLC7A1* #Hs00931450_m1, *ACTB* #Hs01060665_g1) on a CFX96 Thermocycler (Bio-Rad). Samples were analyzed by normalizing expression levels to *ACTB*, and relative quantification was performed using the ΔΔCq method.

### Immunocytochemistry

On Day 1 of subculture, BMEC-like cells were washed twice with DPBS, fixed in 100% ice-cold methanol (Fisher Scientific #A412-1) or 4% paraformaldehyde solution (Thermo Fisher Scientific #AAJ19943K2) for 20 minutes, and washed twice with DPBS. Fixed cells were blocked with DPBS containing 5% donkey serum (Sigma-Aldrich #D9663) and 0.3% Triton-X 100 (Sigma-Aldrich #X100), referred to as PBSDT, at room temperature for 1 hour. Cells were then incubated with primary antibody diluted in PBSDT at 4°C overnight. After primary antibody incubation, cells were washed three times with DPBS, incubated with secondary antibody diluted in PBSDT at room temperature for 2 hours, incubated with 4′,6-diamidino-2-pheny-lindoldihydrochloride (DAPI, Thermo Fisher Scientific #D3571) diluted in DPBS for 10 minutes, and then washed with DPBS three times. Secondary antibody incubation was skipped if primary antibody was conjugated with a fluorophore. Images were acquired using Leica DMi8 microscope and analyzed with Fiji ImageJ. All antibody information is listed in Table S2 and S3.

### Statistical analysis

All data are presented as mean ± standard deviation. One-way ANOVA and the Student’s unpaired t-test were used to determine statistical significance as described in each figure legend.

## Results

### Establishment of leptin transport assay

To quantify leptin transport across the BBB monolayer, we subcultured BMEC-like cells on Transwell filters and used ELISA to measure leptin transport across the cell monolayer (Figure 1A). iPSCs were differentiated into BMEC-like cells according to our previously published protocols, where passive and active BBB properties have been confirmed[18]. To mimic the *in vivo* peripheral-to-CNS transport, leptin was added onto the apical side of Transwell filters, and measurements were performed on medium extracted from the basolateral chamber. Apical leptin concentration was manipulated to reflect the difference in transport capacity under different conditions. All basolateral leptin concentrations increased over time in a concentration-dependent manner (Figure 1B). TEER was measured before and after the experiments to ensure maintenance of barrier integrity, where all TEER values were greater than 900 Ω×cm^2^ (Figure 1C), suggesting that differences in leptin transport were not due to paracellular leakage [49]. Overall, the data demonstrated the suitability of this *in vitro* model for examining leptin transport under different conditions.

### Nuclear receptor agonist library screen

To assess the potential influence of nuclear receptor signaling pathways on leptin transport, a nuclear receptor ligand library containing 73 compounds was screened. We selected a 3 µM dose of leptin and a 30-minute incubation to remain in the linear transport phase (Figure S1). To carry out the screen, 10 µM of ligand was added to both sides of Transwell inserts on Day 0 of subculture and leptin transport was quantified the following day. All compounds were assayed in duplicate, and these data can be found in Table S4.

Seven ligands were selected from primary screening for further validation: 6-Formylindolo [3,2-B] carbazole (an aryl hydrocarbon receptor agonist [50]), troglitazone (a peroxisome proliferator-activated receptor (PPAR)-γ agonist [51]), dexamethasone and cortisone (glucocorticoid receptor agonists [52]), N-oleoylethanolamide (PPARα agonist [53]), 17β-estradiol (estrogen receptor agonist [54]), and pregnenolone (pregnane X receptor agonist [55]). These compounds were selected based on several factors, including their overall effect on leptin transport, their statistical significance in the primary screen, whether they were endogenous or synthetic molecules, and their relevance towards physiological functions [54,56–62]. Of these compounds, Troglitazone yielded a trending increase in leptin transport that was not statistically significant, and all other compounds yielded mean values in line with the negative control. In contrast, 17β-estradiol produced a significant increase in leptin transport when assayed across multiple replicates (∼2-fold increase, p<0.05) (Figure 2A). No overt differences in TEER were noted as a function of small molecule treatment (Figure 2B)

**Figure 2.**
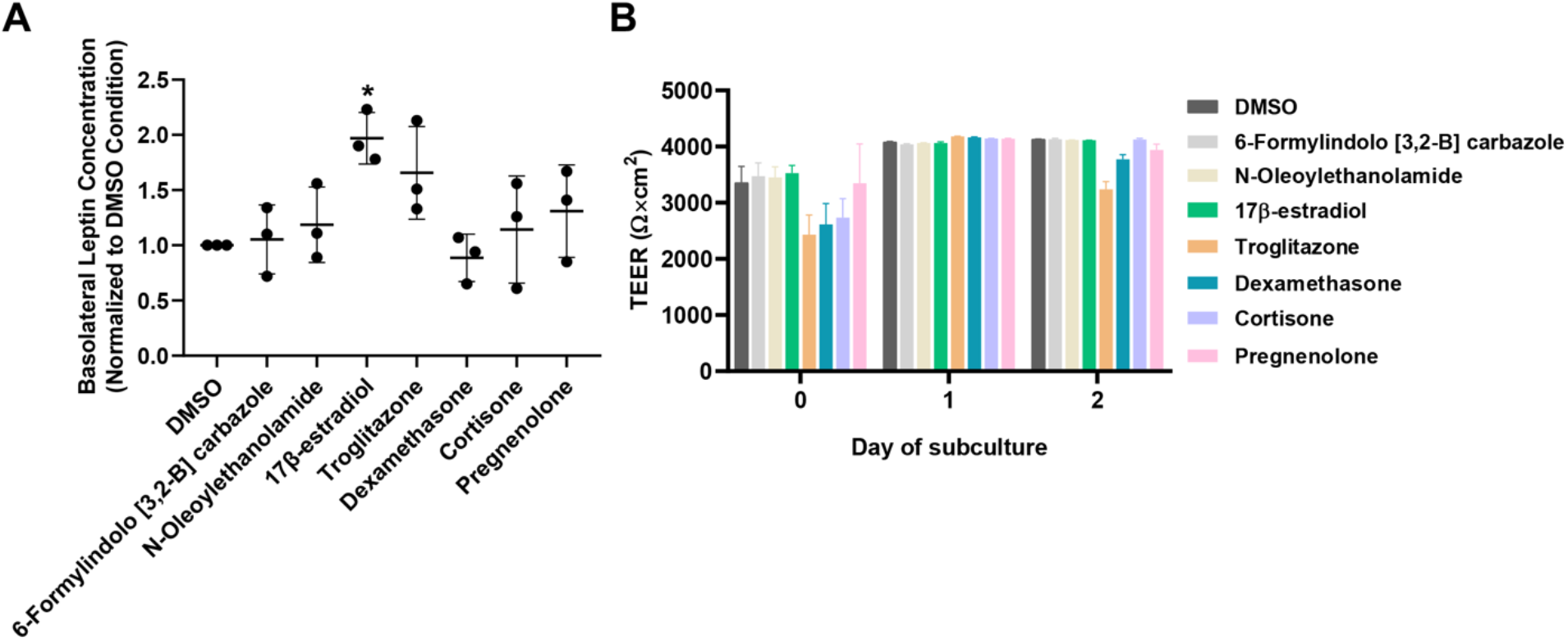
Secondary hit validation for nuclear receptor ligands that regulate leptin transport. For outcomes from the primary screen, refer to Table S4. (A) Compiled datasets represent the basolateral chamber leptin concentration corresponding to the noted ligand treatment, normalized to the DMSO control. Each data point is the mean of a single biological replicate calculated using three Transwell filters (n=3 technical triplicates). Data are represented as mean ± standard deviation from n=3 biological replicates per nuclear receptor ligand. A one-way ANOVA with a Tukey post hoc test was used to evaluate the statistical significance across conditions (*, p<0.05). (B) Representative TEER measurements before and after the leptin transport assay. Data are represented as mean ± standard deviation from 3 Transwell filters (n=3 measurements per filter) per condition used in a single experiment. Trends were confirmed across two additional biological replicates, where no significant differences in TEER were noted as a function of the assayed ligand.

### Transcriptome profiling after 17β-estradiol treatment

Because 17β-estradiol yielded the most pronounced increase in leptin transport, we chose to probe its effects in more depth. To investigate whether 17β-estradiol altered the global transcriptome, RNA was collected from iPSC-derived BMEC-like cells after 17β-estradiol or DMSO treatment and subjected to bulk sequencing (Figure 3A); increased leptin transport by 17β-estradiol treatment and maintenance of passive barrier function was confirmed in these samples (Figure S2). Raw sequencing reads were aligned to the human genome, and differential gene expression was analyzed. Using a threshold of p<0.05, we identified 188 genes as being downregulated and 277 genes as being upregulated within 19,113 genes with non-zero read counts (Figure 3B). Of these, 105 upregulated genes and 57 downregulated genes had a minimum of 1.5-fold change (Figure 3B, Figure S3, and Table S5). To understand similarities between these genes, we employed a Gene Ontology analysis to cluster the enriched gene sets under the 17β-estradiol condition relative to the DMSO condition. From this analysis, we determined that significantly enriched gene sets were predominantly related to transmembrane transporter activity (Figure 3C). Particularly, solute carrier (SLC) genes, including *SLC1A4*, *SLC1A5*, *SLC6A9*, *SLC7A1*, *SLC7A3*, and *SLC7A11*, were recognized under the drug transmembrane transporter activity gene set and found to be significantly upregulated by 17β-estradiol (Figure 3D). We also determined that *LEPR* expression was not significantly altered by 17β-estradiol treatment (Figure 3E), suggesting that changes to leptin transport were not influenced by ObR.

**Figure 3.**
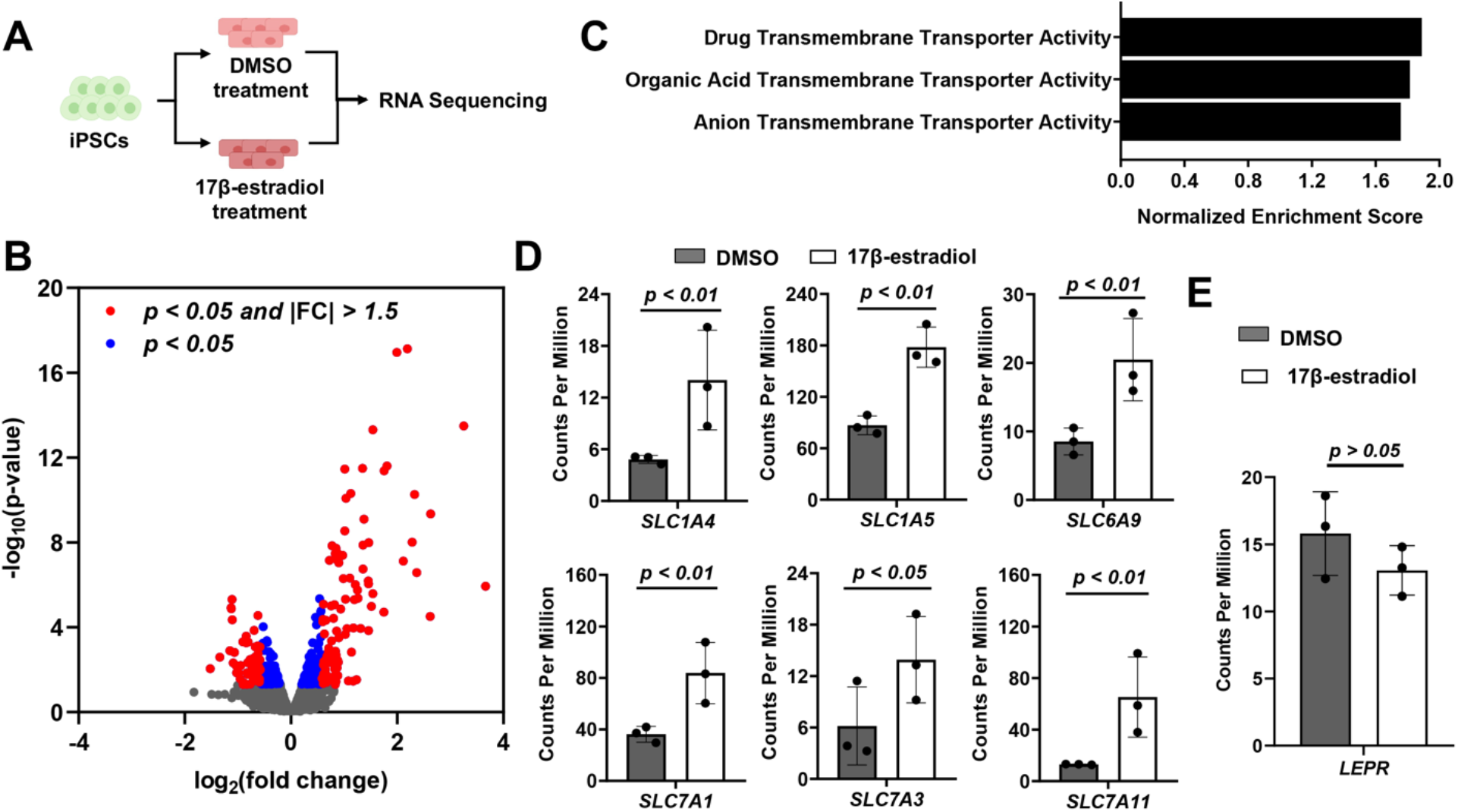
Transcriptome profiling after 17β-estradiol treatment. (A) Experimental setup. 3 biological replicates were sequenced per condition. (B) Volcano plot depicting relative gene expression (17b-estradiol treatment relative to DMSO). Blue dots indicate differentially expressed genes with p<0.05, and red dots indicate differentially expressed genes with p<0.05 and a fold change (FC) greater than 1.5. The full set of differentially expressed genes is provided in Table S5. (C) Gene ontology analysis for enriched signaling pathways (false-discovery rate less than 0.05) in response to 17β-estradiol treatment relative to DMSO. (D) Normalized counts per million for genes encoding solute carrier transporters. Data are represented as mean ± standard deviation from 3 biological replicates.

### Evaluation of SLC transporter contributions to leptin transport

The SLC family is known to facilitate the transport of a range of small molecules, such as glucose, amino acids, and nucleotides [63]. To initially probe links between SLCs and leptin transport, we examined changes to leptin transport in the presence of different amino acids that are predicted substrates of various SLCs. Among these amino acids, substrates of CAT-1 (encoded by *SLC7A1*) augmented leptin transport. Specifically, L-Histidine and L-Ornithine yielded a significant change of ∼1.5-fold (p=0.05) in leptin transport without disruption of passive barrier function (Figure 4). L-Ornithine is also known to be a substrate of CAT-3 (encoded by *SLC7A3*). However, *SLC7A3* has low expression levels in iPSC-derived BMEC-like cells, mouse brain endothelial cells [64], and human endothelial cells [65] compared to the relatively higher expression of *SLC7A1*. In particular, *SLC7A1* is highly and selectively expressed in brain endothelial cells with respect to other neurovascular cell types in single-cell RNA-sequencing datasets [64], marking it as an interesting candidate for further evaluation.

**Figure 4.**
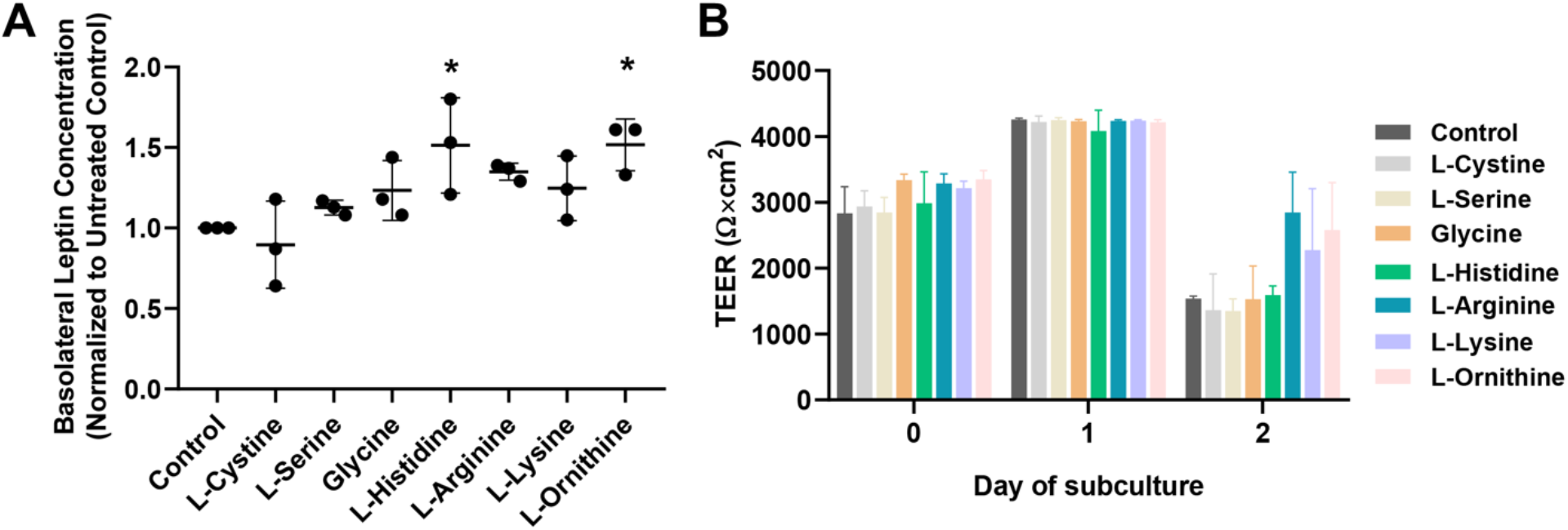
Leptin transport after treatment with different amino acids. (A) Compiled datasets represent the basolateral chamber leptin concentration corresponding to the noted amino acid treatment, normalized to the untreated control. Each data point is the mean of a single biological replicate calculated using three Transwell filters (n=3 technical triplicates). Data are represented as mean ± standard deviation from n=3 biological replicates per amino acid. A one-way ANOVA with a Tukey post hoc test was used to evaluate the statistical significance across conditions (*, p<0.05). (B) Representative TEER measurements before and after the leptin transport assay. Data are represented as mean ± standard deviation from 3 Transwell filters (n=3 measurements per filter) per condition used in a single experiment. Trends were confirmed across two additional biological replicates, where no significant differences in TEER were noted as a function of the assayed ligand.

To more precisely interrogate the role of CAT-1 in leptin transport through modulation of gene expression, an inducible CRISPR platform was introduced into iPSCs with the SB transposon system[44, 48]. We chose to use the CRISPRoff construct which can program heritable epigenetic memory that persists through the differentiation of iPSCs [45]. After transient transfections with plasmids encoding the CRISPRoff system, sgRNAs, and the SB transposase (Figure 5A), iPSCs were subjected to doxycycline treatment to initiate gene silencing through induction of the CRISPRoff expression, then sorted based on BFP expression (Figure 5B). After 9 days of treatment, we confirmed CRISPRoff expression in iPSCs via western blot (Figure 5C), as well as a significant decrease in CAT-1 protein expression (Figure 5D). After differentiating iPSCs into BMEC-like cells, maintenance of the knockdown was confirmed using qPCR to measure *SLC7A1* expression (Figure 5E). In these same cells, leptin transport was significantly increased by ∼2.4-fold relative to the non-targeting sgRNA control cells (Figure 5F). No overt differences in TEER or changes to endothelial or BBB markers were observed after the incorporation of the CRISPRoff system or due to *SLC7A1* knockdown (Figure 5G-H), which strongly suggests that the changes to leptin transport are due to altered CAT-1 expression. Hence, CAT-1 expression and/or activity levels modulate leptin transport within this *in vitro* model.

**Figure 5.**
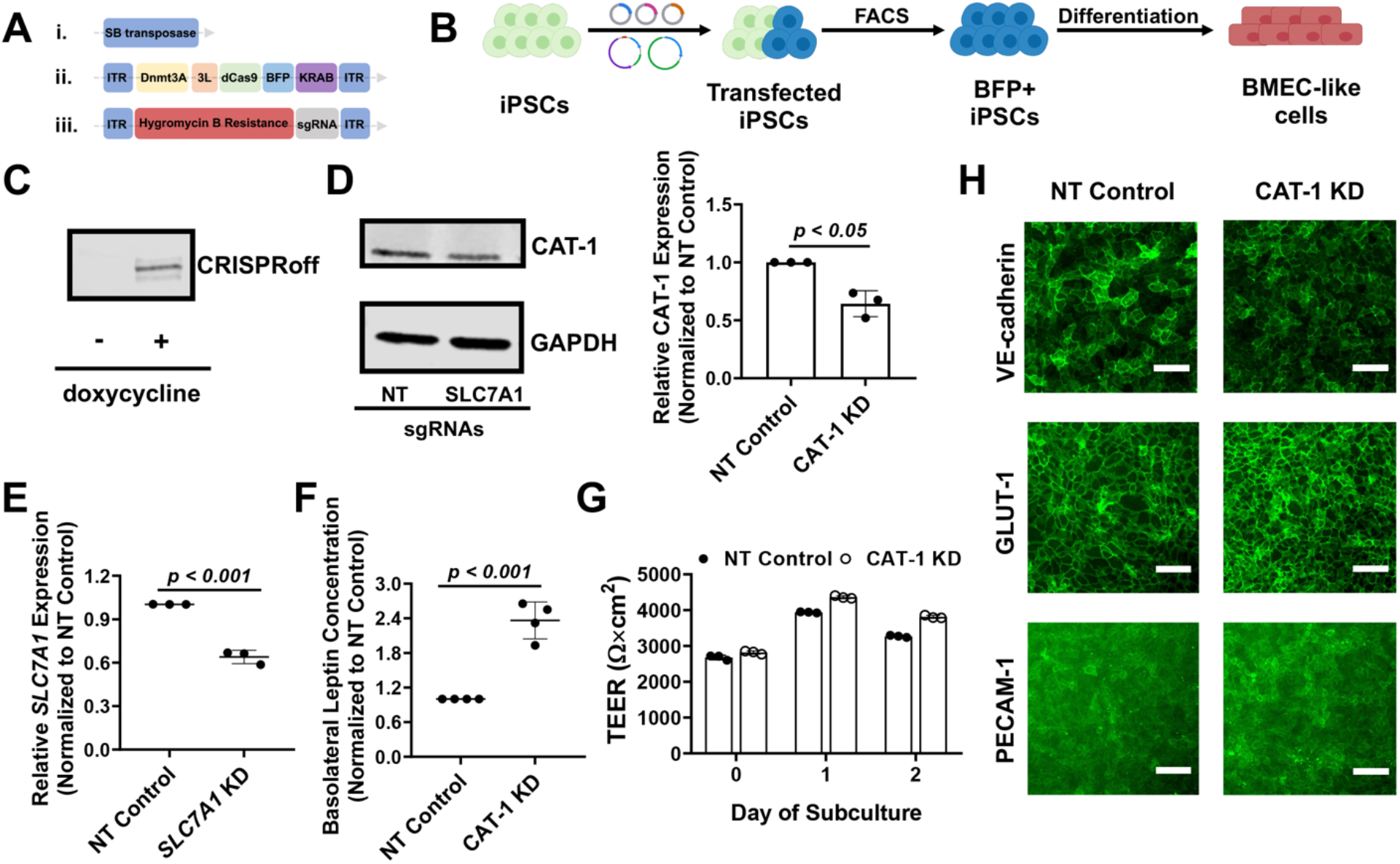
Leptin transport after knockdown of CAT-1 expression. In select panels, NT denotes non-targeting and KD denotes knockdown. (A) Schematic of vectors that were used for generating gene silencing, including SB transposase vector (i), SB transposon/CRISPRoff vector (ii), and SB transposon/U6-sgRNA vector (iii). 3L denotes Dnmt3L. (B) Schematic procedure for iPSC transfection, sorting, and differentiation. After transfections, iPSCs were sorted based on BFP expression. BFP+ cells were then differentiated into BMEC-like cells for the leptin transport assay. (C) Western blot for assessment of dCas9-based CRISPRoff protein in iPSCs with or without doxycycline treatment for 9 days. (D) Western blot for assessment of CAT-1 protein in iPSCs that received a non-targeting gRNA or gRNAs targeting *SLC7A1*. GAPDH served as the loading control. A representative image is shown, and data were quantified using 3 biological replicates (represented as mean ± standard deviation). Each sample was normalized internally to GAPDH and then to the non-targeting control. Statistical significance was calculated using the Student’s unpaired t-test. All iPSC cultures received doxycycline treatment for 9 days prior to sample collection. (E) qPCR assessment of *SLC7A1* expression in BMEC-like cells differentiated from iPSCs that received a non-targeting gRNA or gRNAs targeting *SLC7A1*. Data are represented as mean ± standard deviation from 3 biological replicates. Statistical significance was calculated using the Student’s unpaired t-test. (F) Compiled datasets represent the basolateral chamber leptin concentration corresponding to each gRNA condition, normalized to the non-targeting control. Each data point is the mean of a single biological replicate calculated using three Transwell filters (n=3 technical triplicates). Data are represented as mean ± standard deviation from n=4 biological replicates. Statistical significance was calculated using the Student’s unpaired t-test. (G) Representative TEER measurements before and after the leptin transport assay. Each data point is the mean from a single Transwell filter (n=3 technical measurements per filter). Data are represented as mean ± standard deviation from 3 filters per condition. Trends were confirmed across three additional biological replicates. (H) Representative immunostaining of endothelial or BBB markers in BMEC-like cells in both control and CAT-1 knockdown conditions. Scale bars indicate 100 μm.

## Discussion

The concept of leptin resistance was proposed several decades ago and continues to be explored for its role in obesity disorders. Other than the BBB [8], leptin can penetrate the brain through different routes including the blood-cerebrospinal fluid (CSF) barrier formed by epithelial cells in the choroid plexus [9], the median eminence-CSF barrier composed of tanycytes in the mediobasal hypothalamus [66], and regions allowing the free access to the brain adjacent to circumventricular organs. However, impaired leptin BBB transport has been demonstrated in obese subjects and contributes to functional and behavioral changes related to obesity [1,67–69]. Because the mechanisms of leptin transport across the BBB are mostly unknown, it has been challenging to develop strategies to counteract leptin resistance.

To prospectively identify potential contributors to leptin transport, we combined nuclear receptor ligand screening with transcriptional profiling in our representative *in vitro* BBB model. We first determined that treatment with 17β-estradiol could significantly increase leptin transport across iPSC-derived BMEC-like cells. 17β-estradiol is the predominant circulating estrogen hormone in pre-menopausal women [70], and the circulating 17β-estradiol pool receives contributions from adipose tissues in both males and females [71]. Studies have shown that 17β-estradiol levels are correlated with the incidence of obesity [72], and the use of 17β-estradiol can alleviate the high-fat diet and ovariectomy-induced obesity [61, 73]. The literature regarding the influence of 17β-estradiol on BBB function is very mature, as 17β-estradiol treatment has been shown to regulate the expression of BBB transporters [74–76], BBB permeability, and tight junction integrity [77–80]. However, to our knowledge, no studies have examined global transcriptional changes imparted by 17β-estradiol treatment. Our RNA sequencing and Gene Ontology analyses revealed that 17β-estradiol has a profound influence on transmembrane transporter expression patterns, including the SLC family. We further verified that treatment of BMEC-like cells with SLC substrates, as well as knocking down *SLC7A1*, could significantly alter leptin transport. Thus, while our investigations have not identified a new leptin transporter (since leptin is unlikely to be a substrate for SLC transporters), these studies provide new insight into mechanisms of leptin transport regulation.

A key remaining question is how an amino acid transporter might regulate transport of a peptide hormone across the BBB. There is a precedence for such effects, as studies from several decades ago showed that peptides could influence the passage of nonpeptide substances across the BBB [81, 82], and vice versa [83]. One potential mechanism is modification of intracellular signaling events. For example, arginine methylation has been identified as a key step in Wnt signaling through GSK3 phosphorylation [84], and Wnt signaling has important roles in BBB development [85–87] and maintenance [88]. Since arginine is a substrate of CAT-1 [89], a reduction in CAT-1 expression could impact Wnt signaling activity and yield downstream changes to expression or activity of other transporters; this type of mechanism may also be conserved through other amino acid transporters. Arginine is also the biological precursor for nitric oxide, and it is possible that altered CAT-1 expression could reduce bioavailability of nitric oxide, thereby contributing indirectly to endothelial dysfunction and altered BBB properties [90]. Another potential mechanism is through altered intracellular trafficking. For example, SLC38A9 is essential for an amino acid-stimulated endosome-to-Golgi trafficking in cultured cells through direct interactions with proteins in a GTPase network [91]. In this context, it is possible that altered expression of amino acid transporters changes trafficking within the BBB and enhances transcytosis. Each of these possibilities would require extensive molecular interrogation and will be the subject of future investigations.

## Conclusions

The workflow established in the iPSC-derived BBB model provided a platform to quantitatively assess molecular transport of leptin. Using a nuclear receptor ligand screen, 17β-estradiol was identified for its ability to enhance leptin transport. RNA sequencing revealed bulk transcriptomic changes centered around transmembrane transporters that were differentially regulated by 17β-estradiol. The functional importance of a particular transporter, CAT-1, was further validated through the CRISPRoff system to probe the BBB transport biology. Overall, this work provides important insights on regulation of peptide hormone transport across the BBB and offers a framework for studying the transport of other hormones and nutrients across the BBB using iPSC-based models.

## Abbreviations

BBB: blood-brain barrier
BMEC: brain microvascular endothelial cell
ObR: leptin receptor
TEER: trans-endothelial electrical resistance
iPSC: induced pluripotent stem cell
CNS: central nervous system
ELISA: enzyme-linked immune sorbent assay
CAT-1: cationic amino acid transporter-1
hESFM: human endothelial serum-free media
bFGF: basic fibroblast growth factor
RA: retinoic acid
ECM: extracellular matrix
DMSO: Dimethyl sulfoxide
WebGestalt: WEB-based Gene SeT AnaLysis Toolkit
SB transposon: Sleeping Beauty transposon
BFP: blue fluorescent protein
TRE: tetracycline response element promoter
sgRNA: single guide RNA
DAPI: 4′,6-diamidino-2-pheny-lindoldihydrochloride
PPAR: peroxisome proliferator-activated receptor
SLC: solute carrier
CSF: cerebrospinal fluid

## Declarations

### Ethics approval and consent to participate

Not applicable.

### Consent for publication

Not applicable.

### Authors’ Contributions

YS and ESL designed the research, analyzed the results, and wrote the paper. HK contributed to the RNA sequencing analysis and vector cloning. CAH and JMB provided vectors and advised on implementation of the CRISPRoff system. EMR provided important insight into experimental outcomes. All authors read and approved the final manuscript.

## Acknowledgements

The authors would like to thank Dr. Jamey Young for the use of the BioTek Synergy H1 multi-mode microplate reader. The authors also would like to thank Vanderbilt Flow Cytometry core for technical assistance on cell sorting. Some figures in this manuscript were produced in part with BioRender software.

## Availability of Data and Materials

The datasets used and/or analyzed during the current study are available from the corresponding author upon reasonable request. RNA sequencing data have been uploaded to ArrayExpress under the accession number E-MTAB-11691.

