## Supplemental information for "Nuclear receptor ligand screening in an iPSC-derived *in vitro* blood-brain barrier model identifies new contributors to leptin transport"

**Table S1. Sequences for chimeric gRNAs.**

| Gene Target | Sequences |
|---|---|
| SLC7A1 | GGCACTCACCAAGCCGGCGG |
| SLC7A1 | GATCGCGAACAGACAGACTGT |
| SLC7A1 | GGAGCAGCGGGTCAACTACC |
| Non-targeting control | GATCGTCGGTCGGGTCATACG |
| Non-targeting control | GTTCGTCGCGATAGATGATCG |
| Non-targeting control | GCGCGTATCGTTTCATCGGCG |

**Table S2. Primary antibodies.**

| Target Antigen | Species | Vendor | Product number | Dilution |
|---|---|---|---|---|
| VE-cadherin | Goat | R&D Systems | AF938 | 1:100 |
| GLUT-1 | Mouse | Abcam | Ab195359 | 1:100 |
| PECAM-1 | Rabbit | Thermo Fisher Scientific | RB-10333-P | 1:25 |
| CAT-1 | Rabbit | Proteintech | 14195-1-AP | 1:1000 |
| GAPDH | Mouse | Cell Signaling Technology | D4C6R | 1:1000 |
| Cas9 | Rabbit | Abcam | Ab204448 | 1:1000 |

**Table S3. Secondary antibodies**.

| Species Reactivity | Host | Conjugate | Vendor | Dilution |
|---|---|---|---|---|
| Mouse | Donkey | Alexa Fluor 488 | Thermo Fisher Scientific | 1:200 |
| Rabbit | Donkey | Alexa Fluor 488 | Thermo Fisher Scientific | 1:200 |
| Goat | Donkey | Alexa Fluor 488 | Thermo Fisher Scientific | 1:200 |
| Goat | Donkey | Texas Red | Thermo Fisher Scientific | 1:200 |
| Rabbit | Goat | IRDye 800CW | LI-COR | 1:15,000 |
| Mouse | Goat | IRDye 800CW | LI-COR | 1:15,000 |

**Table S4. Nuclear receptor ligand library primary screen outcomes.** Each compound was assayed in technical duplicate (replicates 1 and 2). Leptin concentrations in the basolateral chambers are normalized to a DMSO control within the same plate to avoid inter-plate variability. Compounds selected for hit validation are colored in blue.

| Ligand | Replicate 1 | Replicate 2 | Adjusted P-value |
|---|---|---|---|
| 25-Hydroxyvitamin D3 | 1.86 | 1.20 | 0.8799 |
| Retinoic acid, all trans | 0.11 | 1.23 | 0.9799 |
| 9-cis Retinoic acid | 0.93 | 1.69 | 0.9851 |
| 13-cis Retinoic acid | 0.27 | 0.06 | 0.5881 |
| 4-Hydroxyphenylretinamide | 2.58 | 3.30 | 0.0551 |
| AM-580 | 1.27 | 1.09 | 0.9203 |
| TTNPB | 1.38 | 1.15 | 0.733 |
| Methoprene acid | 0.82 | 1.16 | >0.9999 |
| WY-14643 | 1.23 | 0.79 | >0.9999 |
| Ciglitazone | 0.52 | 0.41 | 0.7283 |
| Tetradecylthioacetic acid | 4.77 | 0.87 | 0.8999 |
| 5,8,11,14-Eicosatetraynoic acid | 3.04 | 0.52 | 0.9971 |
| **6-Formylindolo [3,2-B] carbazole** | **2.66** | **6.28** | **0.4695** |
| Diindolylmethane | 1.00 | 2.42 | 0.9981 |
| Acetyl-S-farnesyl-L-cysteine | 2.48 | 3.93 | 0.8143 |
| S-Farnesyl-L-cysteine methyl ester | 0.83 | 0.99 | 0.9998 |
| N-Acetyl-S-geranygeranyl-L-cysteine | 0.12 | 1.33 | 0.9721 |
| N-Acetyl-S-geranyl-L-cysteine | 0.74 | 0.78 | 0.9843 |
| Farnesylthioacetic acid | 0.44 | 1.58 | >0.9999 |

| | | | |
|---|---|---|---|
| Bezafibrate | 0.15 | 0.96 | 0.9992 |
| LY 171883 | 1.35 | 1.65 | 0.3587 |
| 15-Deoxy-D12,14-prostaglandin J2 | 0.91 | 0.26 | 0.5188 |
| **Troglitazone** | **0.10** | **0.27** | **0.0782** |
| CITCO | 0.49 | 0.46 | 0.3195 |
| Paxilline | 2.49 | 2.35 | 0.0061 |
| 24(S)-Hydroxycholesterol | 0.01 | 4.02 | 0.9668 |
| 24(S),25-Epoxycholesterol | 0.00 | 1.24 | 0.9996 |
| Pregnenolone-16(alpha)-carbonitrile | 1.28 | 2.75 | 0.9663 |
| Clofibric acid | 0.43 | 0.76 | 0.98 |
| BADGE | 0.32 | 1.77 | >0.9999 |
| GW 9662 | 0.62 | 1.42 | >0.9999 |
| Gemfibrozil | 0.83 | 0.70 | 0.9989 |
| GW 7647 | 0.70 | 2.44 | 0.9449 |
| 3,5-Diiodo-L-thyronine | 0.36 | 1.12 | 0.9982 |
| 3,5-Diiodo-L-tyrosine | 1.15 | 1.81 | 0.9721 |
| Retinyl acetate | 0.11 | 0.60 | 0.8795 |
| 3,5-Diiodo-4-hydroxyphenylpropionic acid | 1.25 | 0.08 | 0.9908 |
| Cholic acid | 0.78 | 0.00 | 0.9008 |
| Deoxycholic acid | 0.96 | 0.79 | 0.7919 |
| Chenodeoxycholic acid | 0.94 | 0.72 | 0.5668 |
| Glycocholic acid | 0.74 | 0.23 | 0.8665 |
| Glycodeoxycholic acid | 0.92 | 0.76 | 0.999 |
| Taurocholic acid | 0.47 | 1.57 | >0.9999 |
| Taurodeoxycholic acid | 0.57 | 0.80 | 0.9791 |
| Rifampicin | 1.10 | 1.97 | 0.8492 |
| **Dexamethasone** | **0.37** | **0.39** | **0.006** |
| Lithocholic acid | 0.21 | 0.02 | 0.0009 |
| 5b-Pregnan-3,20-dione | 0.27 | 0.24 | 0.0022 |
| Adapalene | 1.94 | 1.92 | 0.0559 |
| Farnesol | 1.10 | 0.81 | >0.9999 |
| 3a, 5a-Androstenol | 0.34 | 2.94 | 0.9925 |
| 3α, 5α-Androstanol | 0.38 | 3.33 | 0.974 |
| Z-Guggulsterone | 0.36 | 1.02 | 0.9998 |
| TCPOBOP | 0.97 | 1.79 | 0.9993 |
| **N-Oleoylethanolamide** | **2.91** | **2.60** | **0.7014** |
| GW4064 | 0.87 | 1.26 | 0.9995 |
| Geranylgeraniol | 0.74 | 1.27 | >0.9999 |
| 6a-Fluorotestosterone | 1.72 | 1.44 | 0.2796 |
| Tamoxifen Citrate | 0.58 | 0.70 | 0.838 |
| Mifepristone | 1.15 | 0.28 | 0.9256 |
| Estrone | 0.43 | 1.26 | 0.9973 |
| 13(S)-Hydroxy-9Z,11E-octadecadienoic acid | 1.17 | 1.04 | 0.9996 |
| **Cortisone** | **0.20** | **0.19** | **0.3523** |
| Progesterone | 1.07 | 1.40 | 0.9825 |
| **17β-Estradiol** | **2.23** | **3.04** | **0.0299** |
| **Pregnenolone** | **0.11** | **0.20** | **0.1933** |
| Androstenedione | 0.72 | 0.33 | 0.6533 |
| 1a,25-Dihydroxyvitamin D3 | 0.14 | 0.37 | 0.2779 |
| Docosa-4Z,7Z,10Z,13Z,16Z,19Z-hexaenoic acid | 0.90 | 0.75 | 0.9995 |
| 3-Methylcholanthrene | 0.88 | 1.92 | 0.9762 |
| Acitretin | 0.09 | 0.96 | 0.9522 |
| Pioglitazone HCl | 0.57 | 0.48 | 0.9502 |
| 4-Hydroxyretinoic acid | 0.51 | 0.32 | 0.8969 |

**Table S5. Differentially expressed genes and between 17β-estradiol- and DMSO-treated BMEC-like cells.** Data contained in separate spreadsheet. Gene ontology groupings are also provided.

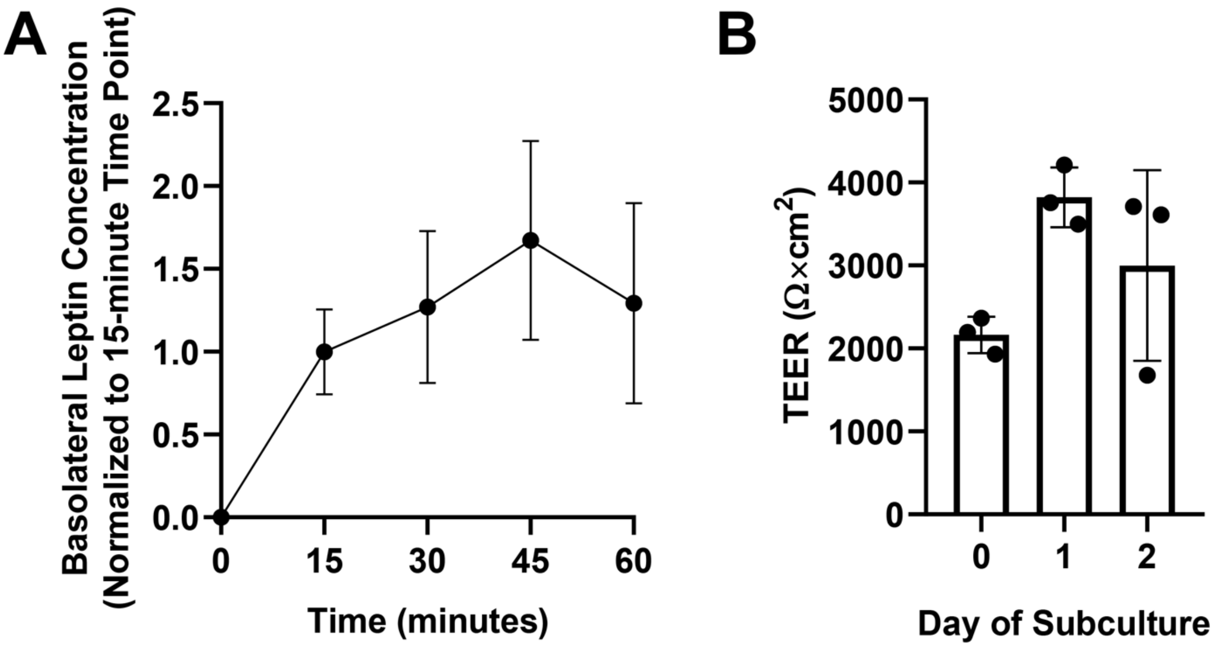


**Figure S1. Assessment of leptin basolateral concentration over time.**

(A) Basolateral leptin concentration was examined every 15 minutes for a 60-minute period. Initial apical concentration was 3 µM. All samples were normalized to the 15-minute time point. Data are presented as mean ± standard deviation from n=3 biological replicates.

(B) TEER measurements before and after the leptin transport assay. Each data point is the average TEER measurement across 3 locations in a single filter, and data are represented as mean ± standard deviation from 3 filters (n=3 technical replicates, n=1 biological replicate). Trends were confirmed in two additional biological replicates.

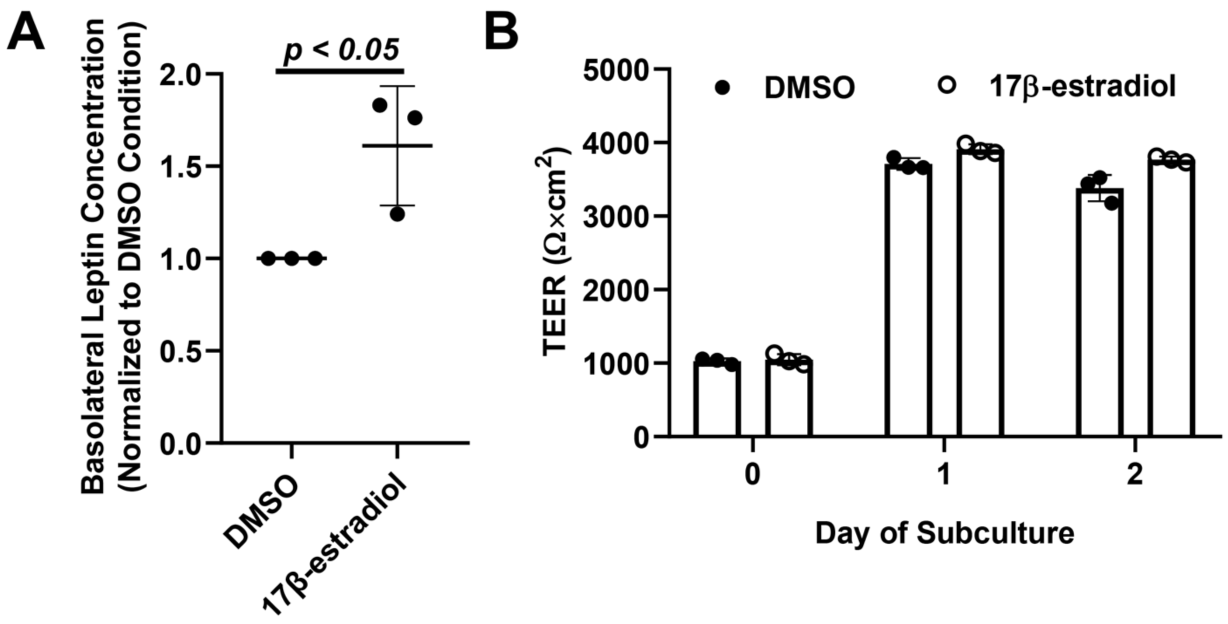


**Figure S2. Leptin transport data corresponding to the samples used for RNA sequencing.**

(A) Compiled datasets represent the basolateral chamber leptin concentration corresponding to 17β-estradiol or DMSO treatment, normalized to the DMSO treatment. Each data point is the mean of a single biological replicate calculated using three Transwell filters (n=3 technical triplicates). Data are represented as mean ± standard deviation from n=3 biological replicates. Statistical significance was calculated using the Student's unpaired t-test.

(B) TEER measurements before and after the leptin transport assay. Each data point is the average TEER measurement across 3 locations in a single filter, and data are represented as mean ± standard deviation from 3 filters (n=3 technical replicates, n=1 biological replicate). Trends were confirmed in two additional biological replicates.

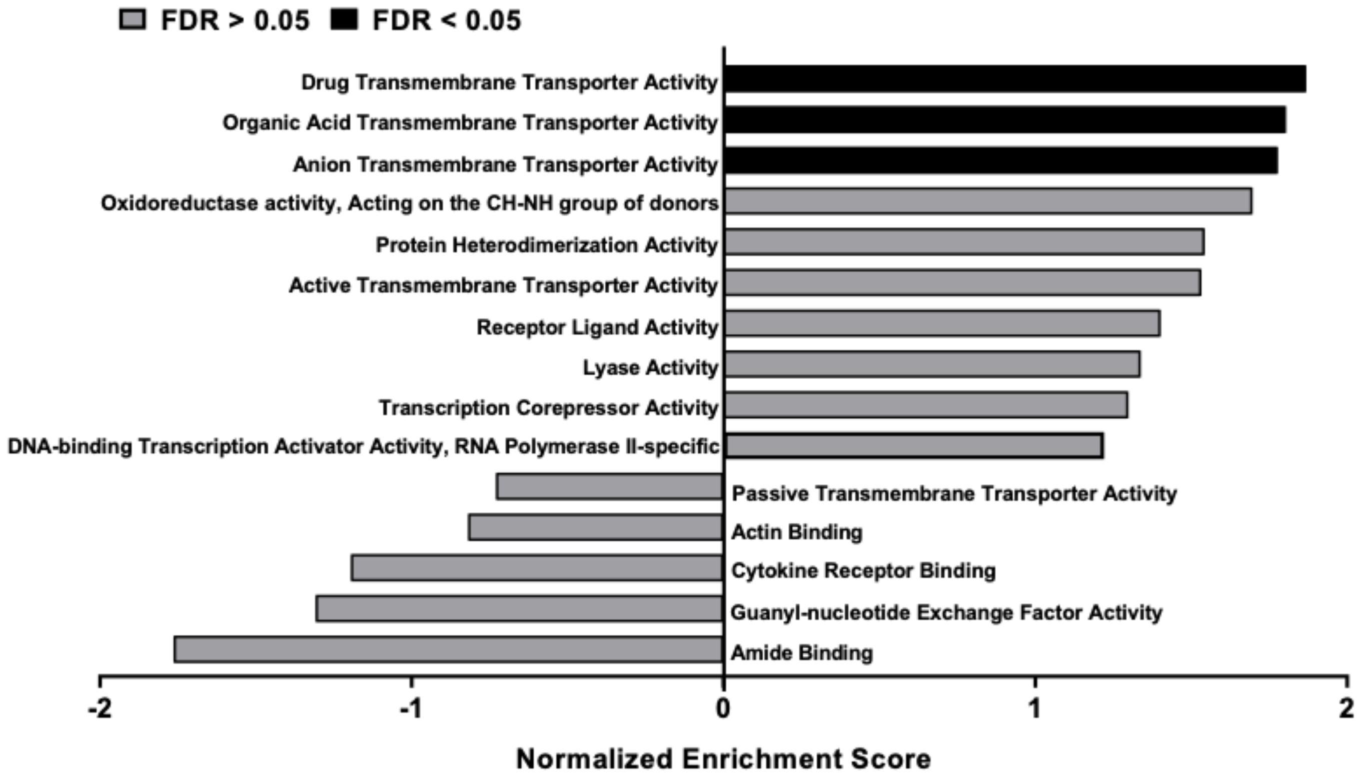


**Figure S3. Gene ontology analysis for enriched signaling pathways in response to 17β-estradiol treatment relative to DMSO.**
